# EDTA tubes are suitable for insulin and C-peptide measurement in resource-limited settings and can be stored at room temperature for up to 24 hours

**DOI:** 10.1101/2024.10.01.616204

**Authors:** Nathan Mubiru, Rogers Mukasa, Isaac Sekitoleko, Priscilla A Balungi, Ronald M. Kakumba, Terry Ongaria, Hubert Nkabura, Moffat Nyirenda, Anxious J Niwaha, Wisdom P Nakanga

## Abstract

**Introduction:** Insulin and C-peptide assessment are important in characterization and management of diabetes. However, their adoption and increased clinical use in low resource settings (LRSs) is partly hindered by logistical factors including supplies required for pre-analytical sample handling and limited infrastructure. We aimed to determine the effects of altered sample processing conditions on stability of insulin and C-peptide at the pre-analytical stage.

**Methods:** We investigated the stability of C-peptide and insulin in serum and plasma collected, preservative type, time to centrifugation, storage conditions and duration of storage on the stability of C-peptide and insulin over 24 hours.

**Results:** Both C-peptide and insulin levels remained stable above 90% from baseline p=1.000 & p=0.776 over 24 hours for samples stored in K2EDTA tubes, whether at room temperature or in a cooler box, both as centrifuged and uncentrifuged whole blood. In contrast, samples collected in plain serum tubes kept at room temperature and uncentrifuged C-peptide and insulin levels decreased significantly to 51%, p=0.006 and 62%, p=0. 083 respectively, similarly insulin levels for centrifuged samples declined to 64%, p=0.083 All iced and centrifuged serum samples remained above 90% of baseline concentration.

**Conclusions:** In resource limited settings where insulin and c-peptide tests are limited to central laboratories and highly dependent on sample referral systems, these tests can be reliably measured without the need for immediate centrifugation or processing from samples collected in whole blood K_2_EDTA tubes uncentrifuged kept at room temperature and processed within 24hours.

**Message:** K_2_EDTA tubes can be used for sample collection in resource limited setting kept at room temperature for up to 24hrs for insulin and c-peptide assays.

K_2_EDTA plasma is more stable than serum for insulin and c-peptide measurement and should be used in resource limited settings

## Introduction

Currently approximately 422 million people worldwide have diabetes and the majority live in low- and middle-income countries(1, 2). Appropriate diagnosis and monitoring of diabetes in people living in low-and middle-income countries is essential and can help improve the quality of life of people with diabetes(3). Evaluation of endogenous insulin secretion, particularly outside the honeymoon period may improve diabetes management in cases where decisions regarding treatment with insulin injections have to be made (4). This can be done by measuring circulating insulin in peripheral blood or C-peptide - a byproduct of proinsulin co-secreted in equal quantities as insulin(3, 5-7).

Of the two, C-peptide is preferred because unlike insulin which undergoes significant hepatic extraction, C-peptide does not undergo first-pass metabolism(5, 8). Recent advances in test methods have made testing of insulin secretion using C-peptide less expensive, more reliable and widely available even in low resource settings. Though the clinical utility of C-peptide and insulin measurement is established, their clinical use is limited by the fragility of both C-peptide and Insulin. These two molecules especially the latter are rapidly degraded both in-vivo and in-vitro by peptidase activity in the blood(9, 10). As a result, the manufacturers of these assays recommend complex pre-analytical sample handling(11, 12). The precarious requirements in the pre-analytical phase may potentially limit its utility particularly in low resource settings were such tests are limited to central laboratories. This implies that sample collection and laboratory analysis happens in different settings, most times hundreds of kilometers apart(13). The standard pre-analytical recommendations involving immediate centrifugation and refrigeration may not be feasible in the periphery settings and/or diabetes clinics(14). In the field of clinical diagnostics, the stability of analytes during the pre-analytical phase is of paramount importance(15) however, in resource-limited settings where access to sophisticated laboratory equipment and infrastructure is limited, preserving the integrity of blood samples becomes a critical concern(16).. McDonald et al and other researchers have previously shown that C-peptide and insulin are stable in dipotassium ethylene diaminetetraacetic acid (K2-EDTA) whole blood tubes at room temperature for up to 24 hours(17, 18). However, it is unclear as to whether these findings are applicable in the tropical areas of SSA were temperatures are higher.

We aimed to determine the most appropriate sample collection and processing criteria for insulin and C-peptide applicable in re source-limited settings by investigating stability over 24hours at variable conditions.

## METHODS

### Ethical clearance

We conducted this study at the Clinical Diagnostics Laboratories within the Medical Research Council/Uganda Virus Research Institute and London School of Hygiene & Tropical Medicine Uganda Research Unit. Written informed consent was obtained from all participants before participating in the study. The study was approved by the Uganda Virus Research Institute Research Ethics Committee (ref: GC/127/19/03/700), Uganda National Council for Science and Technology (ref: HS 2566), and London School of Hygiene and Tropical Medicine Ethics Committee (ref: 17514).

### Procedures

#### Subjects: Eligibility criteria

Participants willing to provide informed consent, aged ≥18 years, with no history of diabetes.

After obtaining informed consent, we collected data from participants on; age, sex and fasting status using a structured questionnaire.

Non fasting 40mls whole blood venous samples were collected for insulin and C-peptide measurements using an 18G butterfly needle and 20ml syringe. This was quickly dispensed into 20 Dipotassium Ethylenediamineteraacetic acid (K2EDTA) tubes and 20 plain serum tubes (figure.1).

**Figure 1.**
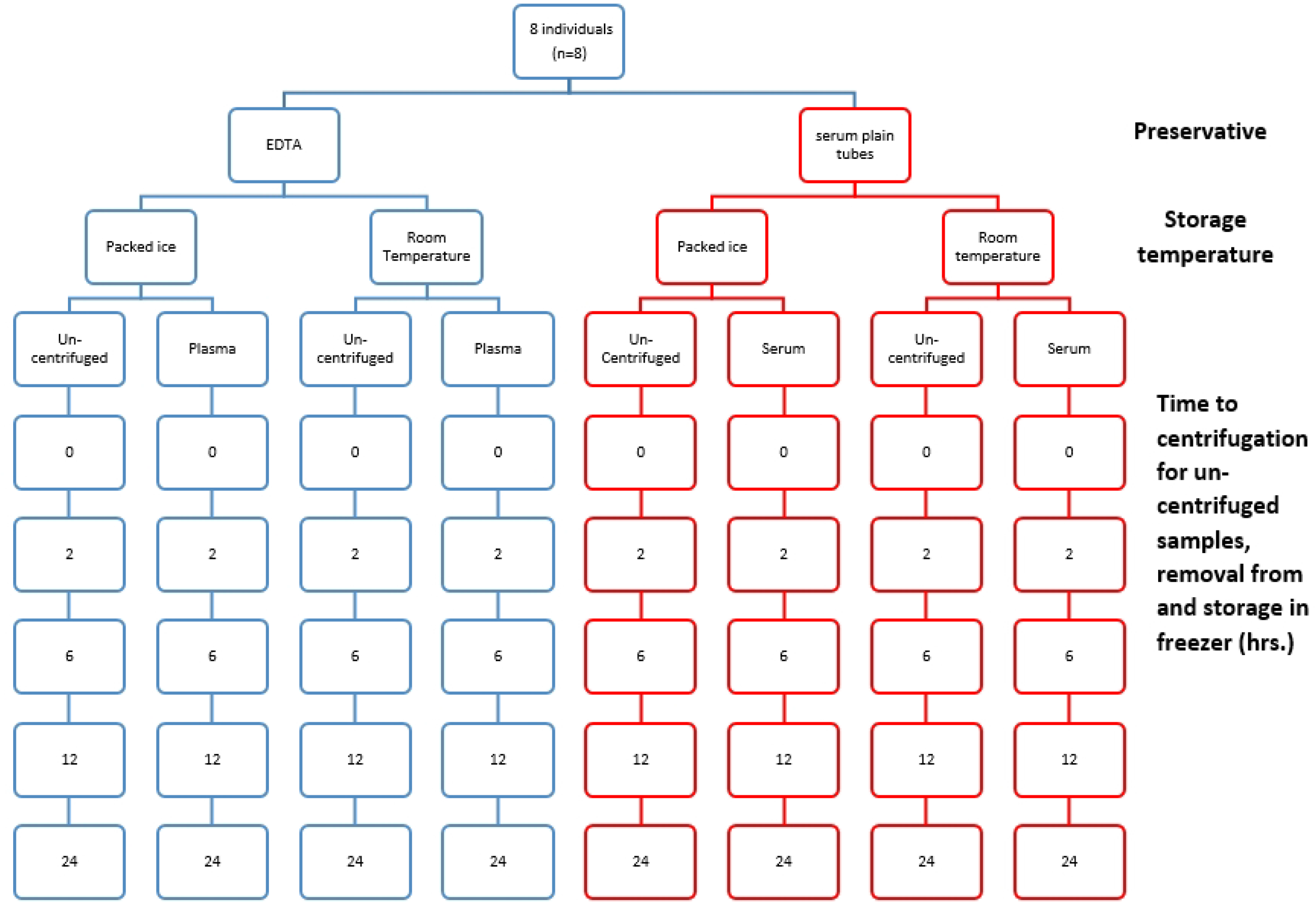
Flow diagram showing sample collection study protocol for stability of c-peptide and insulin in K2EDTA and plain serum tubes over 24 hours. At each time point samples were centrifuged at 3000rpm for 10 minutes and supernatant stored at -20^°^C

At zero time (baseline), four K2EDTA and four plain serum tubes were centrifuged at speed of 3000RPM for 10 minutes, for serum samples these were left to clot 30 minutes after collection prior to centrifugation. Each tube was then divided into 4 aliquots and stored in a -20°C freezer as reference specimens.

At 2, 6, 12, and 24 hours, two K2EDTA and two plain serum tubes were centrifuged at speed of 3000RPM for 10 minutes. Aliquots were prepared and divided. Two aliquots from each tube type were kept at room temperature, while the others were stored in a cool box with ice.

At each time point, one aliquot from each tube stored at room temperature and in the cool box was placed in the -20°C freezer. Additionally, four K2EDTA and four plain serum tubes were stored at room temperature. At 2, 6, 12, and 24 hours after sample collection, one tube of each type was centrifuged, aliquoted, and placed in the -20°C freezer.

Two K2EDTA and two plain tubes were stored in the cool box following the same procedure as those stored at room temperature.

All samples were then frozen at -20^°^C and prior to analysis, all frozen samples were thawed once for 30-60 minutes to reach room temperature (maintained at 18–26°C). These were analyzed as a batch at the MRC/UVRI and LSHTM Clinical Diagnostics Laboratory in Entebbe, Uganda. The Cobas 601 analyzer (Roche/Hitachi, Tokyo, Japan) was used for C-Peptide and insulin measurement using the electrochemiluminescence method. The C-peptide assay uses a direct electrochemiluminescence immunoassay using mouse monoclonal anti-C-peptide antibody labelled with ruthenium and a second mouse monoclonal anti-C-peptide antibody coupled to paramagnetic particles. The insulin assay uses a similar method of two mouse monoclonal anti-insulin antibody coupled to paramagnetic particles.

We compared the results of C-peptide and Insulin samples kept in four different pre-analytic conditions: (1) un-centrifuged kept at room temperature, (2) un-centrifuged kept in a cool box with ice, (3) centrifuged kept at room temperature, (4) centrifuged kept in a cool box with ice.

### Statistical analysis

Participants characteristics were summarized using frequencies and proportions for categorical variables and median (IQR) for the continuous variables. We then computed the mean % change in C-peptide and insulin results from baseline ± 95% confidence interval. Differences between the baseline and the concentration of C-peptide and insulin at 24 hours under the 8 different pre-analytical conditions were examined using Wilcoxon signed rank. Our results were considered clinically significant if the percentage changes of insulin and C-peptide from baseline concentrations was >10% and statistically significant if the p value was <0.05.

## Results

Eight participants (six men and two women, mean age, 30.8 ± 9) with no history of diabetes volunteered to take part in the study. The mean ambient room temperature ranged from 25°C to 28°C and cool box temperatures ranged from, 6°C to 15°C There was no significant difference in the mean baseline insulin levels (864 pmol/L for K2EDTA and 800 pmol/L for serum, P=0.383) or in the mean c-peptide levels (17.3 μU/mL for K2EDTA and 14.7 μU/mL for serum, P=0.312)

### C-peptide and insulin were more stable in samples collected and kept at room temperature in K2EDTA than those collected in plain serum tubes with or without centrifugation

C-peptide and insulin stayed more stable in unprocessed whole blood samples collected in tubes with K2EDTA compared to plain serum tubes. Over 24 hours, their levels in K2EDTA tubes remained above 90% of the initial levels (90% of baseline for C-peptide and 98% for insulin), with no significant decrease (p=1.000 for C-peptide and p=0.404 for insulin). In contrast, C-peptide and insulin levels significantly dropped in plain serum tubes within 24 hours, falling to 51% and 64% of baseline respectively (p<0.001 for both). (Figures 2C and 3C, Figure 2D and 3D)

**Figure 2.**
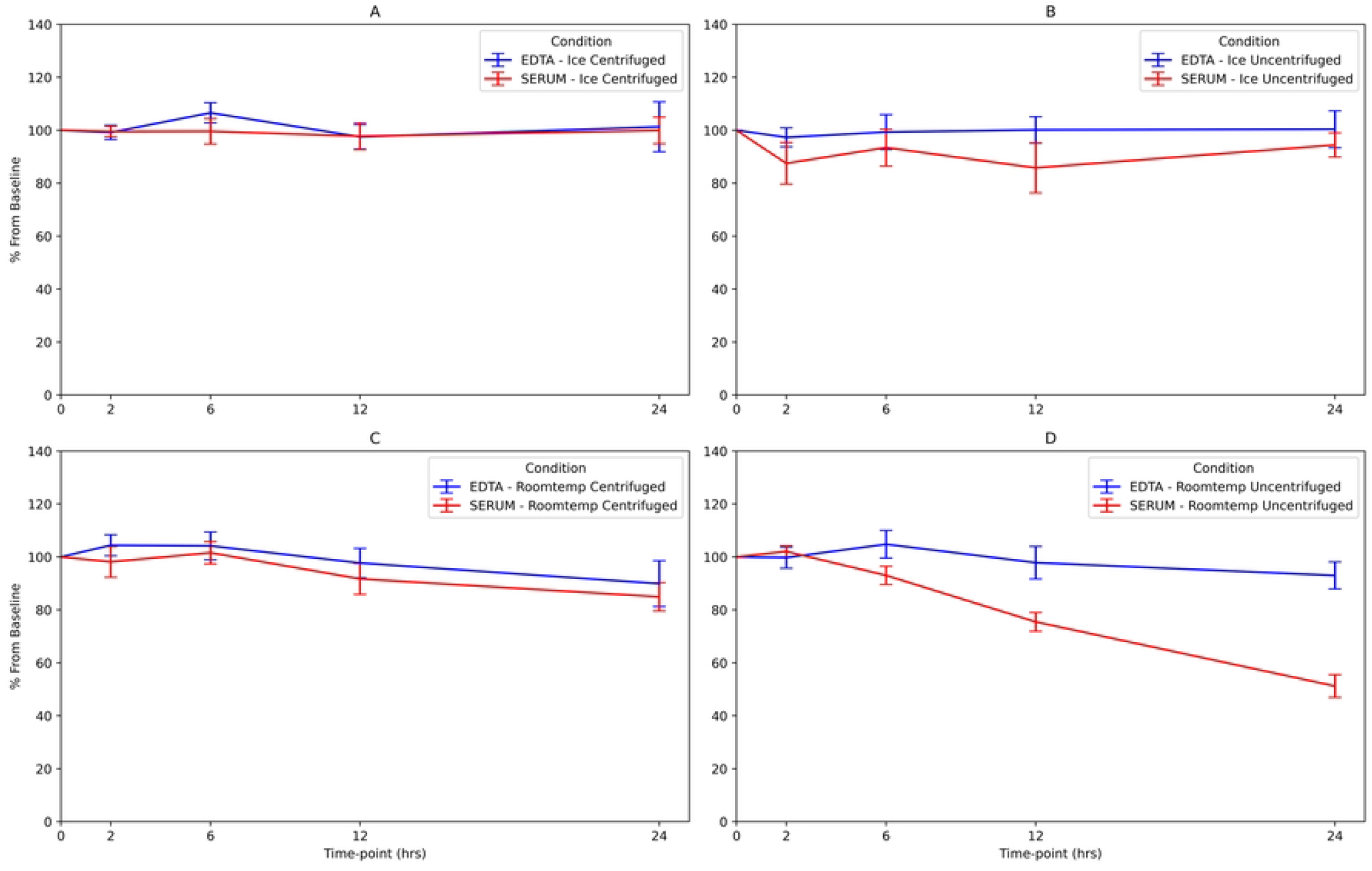
Graphical presentation of % decrease for c-peptide from baseline at different conditions

**Figure 3.**
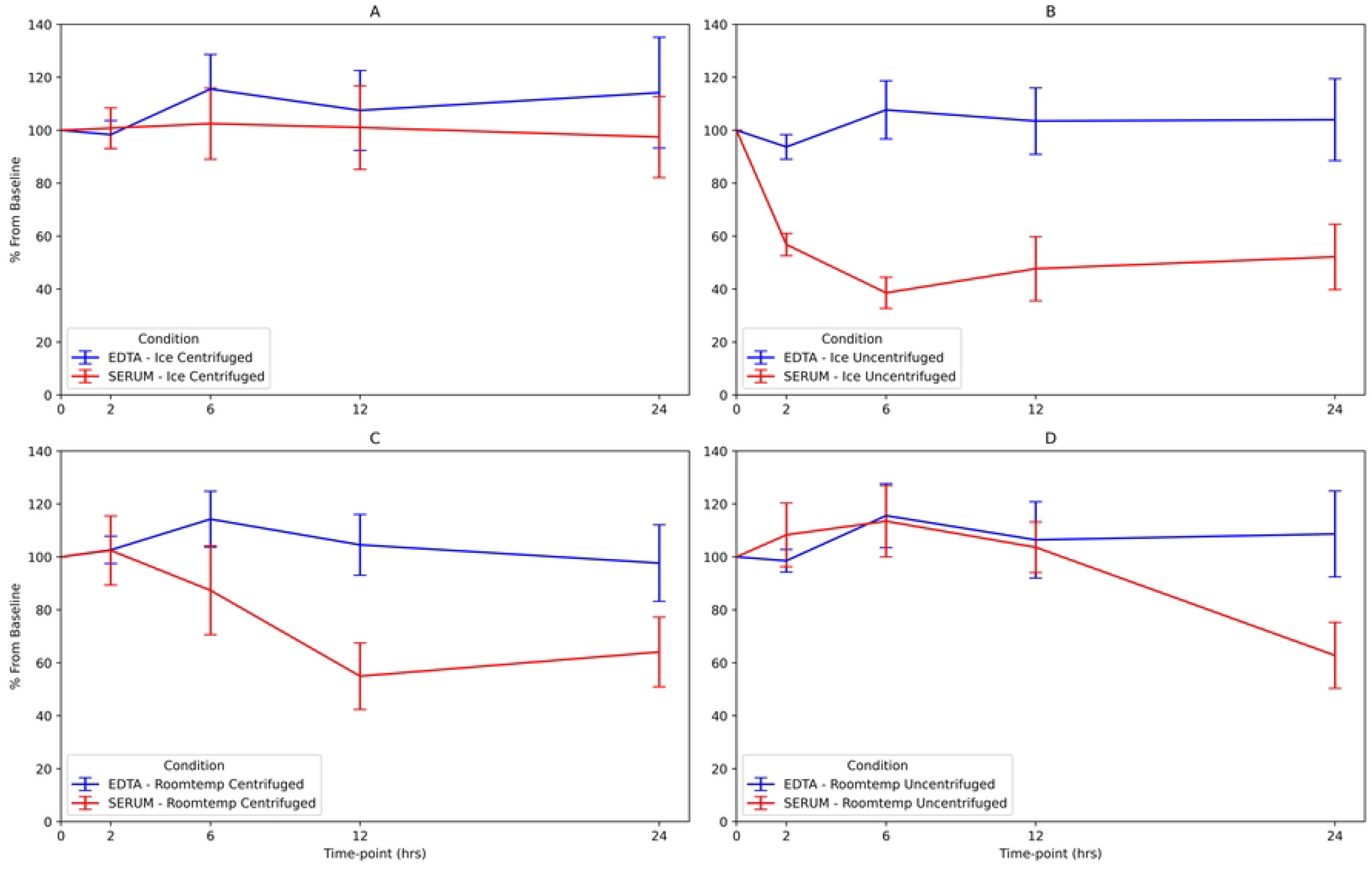
Graphical presentation of % decrease for Insulin from baseline at different conditions

For centrifuged samples, C-peptide in plain serum tubes dropped to 85% (p<0.001), while insulin dropped to 62% (p=0.083). (Figure 2D and 3D)

### C-peptide was stable in K2EDTA tubes and Serum tubes both as centrifuged and uncentrifuged kept in cooler box

C-peptide levels, in both centrifuged and unprocessed whole blood, did not decrease below 90% of baseline (p-1.00) when the samples were kept in a cooler box with ice up to 24 hours (figure 2A and 2B).

### Insulin levels in Centrifuged and un-centrifuged K2EDTA were stable for up to 24hours when kept in a cool box

Insulin levels did not drop below 90% of baseline over 24 hours for samples kept in a cooler box with ice, both as centrifuged and unprocessed whole blood K_2_EDTA (113% of baseline, p=1.000 and 104%, p=0.404 of baseline) (Figure 3A and 3B).

### Insulin levels in serum remained stable when samples were centrifuged, and vice versa for un-centrifuged for samples kept in a cool box

In contrast, insulin concentrations decreased significantly in samples stored in plain serum tubes for uncentrifuged but not for centrifuged samples (52% of baseline, p=0.006 and 97% of baseline, p=1.000 at 24 hours respectively), (Figure 3C and 3D).

### Insulin and c-peptide levels were stable at 0min,2hr, 6hr and 12hr in both plain serum tubes and K2EDTA for most conditions except at highlighted conditions below

C-peptide and insulin samples collected in plain serum tubes, when kept on ice and uncentrifuged, dropped below 90% of baseline within 2 hours. Insulin levels dropped to 50% and c-peptide to 86% of their initial concentrations across all time points up to 24 hours, (Figure 2B and 3B).

C-peptide samples collected in plain serum tubes and left at room temperature without centrifugation decreased to 54% of baseline concentration by 12 hour), (Figure 2D).

### Overall stability of insulin and c-peptide in serum and K2EDTA

K2EDTA tubes improved stability and did not drop below 90% of baseline concentration across all conditions for samples collected aimed at testing insulin and c-peptide as compared to the plain serum tubes.

## DISCUSSION

Our study found that insulin and C-peptide remain stable when collected in K2EDTA tubes un-centrifuged whole blood for at least 24 hours at ambient room temperatures. In addition, we also found that insulin and c-peptide remained stable when samples were collected in K2EDTA and kept in a cool box whether centrifuged or uncentrifuged for up to 24hrs. Our results also showed that insulin and c-peptide remained stable in centrifuged plain serum tubes kept in a cool box for up to 24hrs, for uncentrifuged serum samples these remained stable for up to 2hrs. Furthermore, our findings divulged that serum samples left at room temperature, unprocessed all drop below 90% of baseline concentration by 24hrs. Finally, from our results we revealed that K2EDTA tubes improved stability of insulin and c-peptide and did not drop below 90% of baseline concentration across all conditions as compared to the plain serum tubes within 24hrs.

Studies by McDonald et al (2012) and Déchelotte et al (2016) support these findings showing that insulin and C-peptide are stable in K2EDTA tubes for up to 24hrs.Incontast some assay manufacturers have reported stability of 4 hours at 20-25 °C(19, 20) contrary to our findings. Previous studies documented that insulin and c-peptide samples when centrifuged immediately and kept in a cool box, were all stable regardless of whether they were collected in serum or K2EDTA tubes and are in line with finding from other authors and assay manufactures who reported stability of up to 2days at (2-8)^0^ c.(13, 14, 21).

This stability is likely due to the cool temperatures that slowed down the enzymatic process and the centrifugation which separates red cells from the analyte, preventing the release of proteases through cell lysis as well as the fact that K2EDTA tubes contain Dipotassium Ethylene diamine tetra-acetic acid (K2EDTA), a molecule that chalets co-factors required in the proteolytic activity.

In contrast, findings from Nkuna et al (2023) showed c-peptide when collected in K2EDTA tubes left at room temperature was stable for up to 12hr contrary to our finding which showed K2EDTA improved stability up to 24hr which is supported by McDonald et al (2012) and Déchelotte et al (2016) as well.

This study has several strengths. Firstly, we evaluated the impact of delayed centrifugation which is very relevant for referral samples in regards to resource limited settings where collection sites are mostly far from testing facilities. We assed stability over 24hrs at various storage temperatures considering our tropical settings and examined centrifuging time in relation to the possible times it would take referral samples to reach the testing facility.

Limitations for this study include the absence of samples with low insulin or C-peptide levels, therefore we did not evaluate how such samples would perform in this experiment and samples were tested on only one analytical platform hence where unable to compare the level of agreement between analytical platforms. Lastly, the effect of interfering factors like heamolysis and pharmaceutical compounds was not determined.

The fact that insulin and c-peptide were stable at room temperature for up to 24hrs in unpossessed K2EDTA suggests that in resource limited settings with limited access to electricity and pre-analytical processes, samples for testing C-peptide and insulin can be collected in K2EDTA whole blood tubes and processed after at least 24hours without significant alteration. This finding simplifies sample collection procedures, especially in low resource setting hence the same sample can be used for HbA1c and haemoglobin concentration measurements(22, 23). Combining these results with our previous findings showing glucose stability in K2EDTA tubes, it appears that the same tube can be used for collecting samples for insulin, C-peptide and glucose testing as long as processing occurs within 6 hours and kept in a cool box. (24).

Adoption of hand held centrifuges could provide solutions to the common challenges associated with advocating for immediate blood separation, especially in remote areas where conventional centrifuges and electricity are not readily available (25, 26). Additionally, establishment of point of care instruments that can test for insulin and C-peptide could eliminate the need for expensive pre-analytical activities involved in the sample preparation for these assays(27).

## Conclusion

In conclusion, this study has shown that insulin and C-peptide are stable in K_2_EDTA whole blood, without processing when kept at room temperature for up to 24 hours. This implies that in resource limited settings where insulin and c-peptide tests are limited to central laboratories and highly dependent on sample referral systems, these tests can be reliably measured without the need for immediate processing provided they are tested or processed within 24hours.

## Acknowledgement

We acknowledge the participants that took part in the study and the clinical Diagnostic laboratory staff at the MRC, UVRI and LSHTM that supported the diagnostic activities.

## Author Contributions

Nathan Mubiru: writing original draft, review and editing, Methodology

Rogers Mukasa: Methodology and writing –review and editing

Priscilla Balungi Agather; Methodology, Writing –review and editing

Isaac Sekitoleko; Data analysis, Writing –review and editing

Ronald Makanga Kakumba: Data analysis, Data cleaning, writing –review and editing

Hubert Nkabura; Writing –review and editing

Terry Ongaria; Data analysis, Data cleaning

Anxious Jackson Niwaha: Writing –review and editing

Wisdom Petros Nakanga: Formal analysis, Supervision, Investigation, methodology Writing original draft Writing –review and editing

Moffat J. Nyirenda: Funding Acquisition, Conceptualization Writing and Reviewing

## Data Availability Statement

All the relevant data are within the paper and its Supporting information files.

